# New phylogenetic Markov models for inapplicable morphological characters

**DOI:** 10.1101/2021.04.26.441495

**Authors:** Sergei Tarasov

## Abstract

This paper proposes new Markov models for phylogenetic inference with anatomically dependent (inapplicable) morphological characters. The proposed models can explicitly model an anatomical dependency in which one or several characters are allowed to evolve only within a specific state of the hierarchically upstream character. The new models come up in two main types depending on the type of character hierarchy. The functions for constructing custom character hierarchies are provided in the R package *rphenoscate*. The performance of the new models is assessed using theory and simulations. This paper provides practical recommendations for using the new models in Bayesian phylogenetic inference with *RevBayes*.

## Introduction

Recent publications have revitalized the interest in the long-standing problem of modeling inapplicable characters for phylogenetic analysis with discrete morphological data (Maddison, 1993; Hawkins et al., 1997). They offer new approaches for parsimony (Brazeau et al., 2019; Hopkins and St. John, 2021; Goloboff et al., 2021) and probabilistic methods (Tarasov, 2019). In morphology, character inapplicability refers to the hierarchical dependency of traits due to the anatomical dependency of their respective body parts. For example, tail color is absent (i.e., inapplicable) if a tail is absent but can be blue or red if the tail is present. Evolutionarily this means that the color trait (i.e., the dependent character) evolves only within the certain state of the controlling character when the tail is present. Hereafter, I call this the “embedded dependency” (ED) to distinguish it from other types of dependencies that may exist between characters. Anatomical hierarchy may induce EDs of arbitrary complexity, which may involve several inapplicable characters. For example, a two-level dependency may include color for a tail armor that depends on the armor’s presence; in turn, the armor depends on the tail’s presence. Due to anatomical dependency between body parts, the ED is widespread in morphological data but not solely confined to them; it also occurs in molecular data. For example, if we consider a particular DNA site as the character, its states are the four nucleotides *{A,C,G,T*}. They depend on the gene in which this site is located. If this site is absent due to the birth-death of the gene, then those four states are inapplicable too.

Currently, two probabilistic approaches exist for modeling ED. The multistate stochastic Dollo (MSSD) approach can model ED with a one-level hierarchy that includes one controlling and one or more dependent characters (Alekseyenko et al., 2008). Inspired by Dollo’s law of irreversibility, it assumes that the birth of a controlling trait, for example, a gain of a tail, may occur only once over the entire phylogenetic tree. However, multiple losses of this trait are allowed. The transitions between the states in the dependent character (e.g., tail color) may occur only when the controlling trait is present. MSSD approach is a good match for modeling ED in which the controlling trait is born only once, but it might be too restrictive if the controlling trait undergoes repetitive births over phylogeny. Such a homoplastic evolution is often observed in morphological characters.

Another approach proposes to model ED using structured Markov models (SMMs) in which anatomical dependency is simulated using hidden states (Tarasov, 2019). This approach allows multiple gains and losses of the controlling trait and can model ED corresponding to a complex anatomical hierarchy with several characters. A recent study (Goloboff et al., 2021) criticized the SMM approach for inappropriately modeling the ED by claiming that hidden states do not reflect anatomical dependencies. I agree with Goloboff et al. (2021) that SMMs do not explicitly imply the ED but disagree that they treat ED incorrectly; SMMs can model various dependencies and behave appropriately for the ED, as was shown in the simulations and analytical derivations (Tarasov, 2019).

As discussed above, the available probabilistic models for ED either impose restrictions on the hierarchical complexity and number of births in controlling trait (MSSD) or do not explicitly capture the ED process (SMM). Thus, the phylogenetic community is still missing a general ED model that would be extensible to custom hierarchies where controlling traits are allowed multiple births. This paper proposes a new class of Markov models that explicitly model the ED and fulfill those requirements. They can be constructed using the techniques for character amalgamation and aggregation proposed for SMMs (Tarasov, 2019, 2020). These techniques are central to modeling morphological characters since they equip them with a mathematically consistent closure property – characters can be added together (i.e., amalgamated) to yield another character, and a character can be divided by the state aggregation to yield one or few characters. The provided ED models come in two flavors depending on the type of characters involved in a hierarchy. This paper describes an algorithm for constructing any custom hierarchy with the ED models and gives practical examples of using them in phylogenetic inference with RevBayes (Höhna et al., 2016). The functions for constructing custom hierarchies are implemented in the R package *rphenoscate*. To test the proposed models, I perform simulations using the tail color problem (Maddison, 1993) and its modified version, the “tail color and armor case”, which well-exemplify anatomical dependencies and were investigated earlier with SMMs (Tarasov, 2019). The simulations demonstrate the appropriate behavior of the proposed models. The focus of this paper is morphological characters; however, in the end, I briefly discuss the usage of the proposed models for molecular data.

## Embedded Dependency Process

### Preliminaries

For rigor, I find it necessary to describe the ED process using precise definitions that will be used throughout the paper and help differentiate between the existing models. In characterizing the ED, I follow earlier studies (Sereno, 2007; Dececchi et al., 2015) and distinguish between two main types of morphological characters: a *birth-death (BD) character* that denotes the absence or presence of an anatomical structure (e.g., tail), and a *qualitative character* that describes qualities of a structure (e.g., shape or color). Below, using two motivating examples of tail evolution, I show that these two types may have different evolutionary dynamics.

Due to anatomical dependencies, any ED includes at least one, *controlling character* whose presence implies the presence of a hierarchically downstream character(s), and at least one, *dependent character* whose presence depends on a controlling one. Controlling characters are always of the BD type since only gain or loss (i.e., presence or absence) of such a character can anatomically control the presence or absence of downstream characters. Instead, the dependent characters can be either of the two types. For example, both tail color (qualitative character) and tail armor (BD character) are dependent on the tail’s presence.

### Qualitative ED process (ED-ql)

Let us consider the evolution of tail color in species with no tails, blue tails, and red tails (Maddison, 1993). We are interested in assessing how anatomical dependencies may affect the evolution of those phenotypes. Suppose that the tail evolves through repetitive gains and losses, creating time intervals in a phylogenetic tree when it is present or absent. Clearly, the tail color can only transit between its states “red”, and “blue” when the tail is present, and no color evolution occurs when it is absent. The birth of the tail implies that it immediately appears in red or blue color since color is the tail’s inherent property. One possible scenario of this evolution is shown in Fig. 1A. It is the simplest case of the ED process between the controlling BD character (tail) and the dependent qualitative character (color), where the color’s evolution is embedded within the presence of the tail and is inapplicable otherwise. Hereafter, I refer to it as the “qualitative ED process” (ED-ql) after the type of the dependent character.

**Fig. 1.**
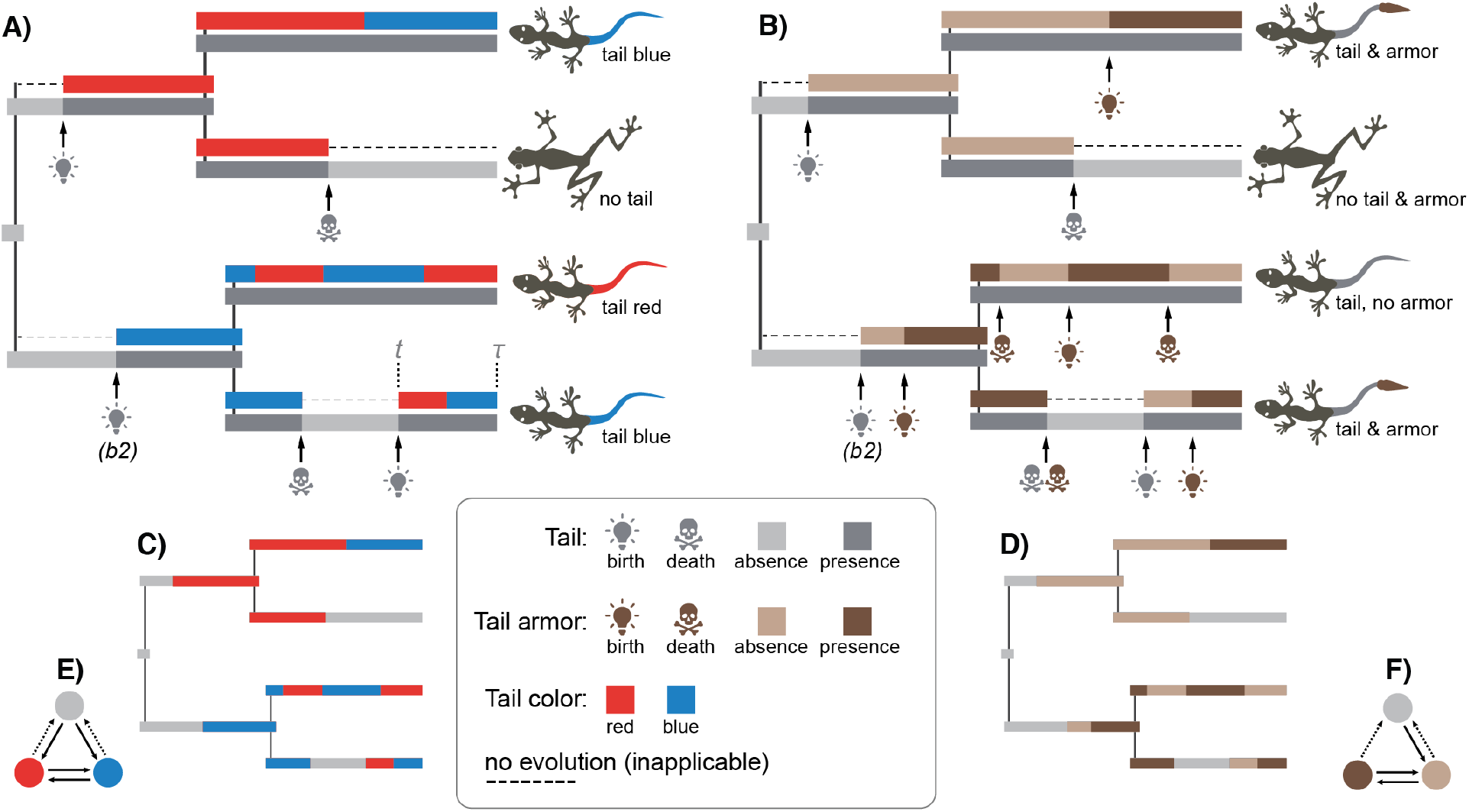
Possible stochastic realizations for the Embedded Dependency (ED) processes. A) The ED-ql process; two characters: tail and color, evolve over phylogeny; the birth-death of the tail generates time intervals when this character is present or absent; the color character (blue or red) only evolves when the tail is present. B) The ED-bd process; two characters: tail and armor, evolve over phylogeny; the tail character evolves as in (A), while the birth-death of the armor character is only allowed when the tail is present; the births of the armor occur with a lag after the births of the tail. C) and D) are the same processes as in (A) and (B), respectively, represented as amalgamated MCs. E) and F) show MCs for the processes in (C) and (D); dashed arrows indicate the same rate parameters. b2 in (A) and (B) indicates the second birth of the tail.

Continuous-time Markov Chain (MC) is the natural choice for modeling discrete characters. Following the logic of Markov processes, one may think that the tail and color evolve as two interacting MCs: one MC characterizes the birth-death evolution of the tail, while another MC, which describes the color character, is allowed to evolve exclusively when the tail is present.

### Birth-death ED process (ED-bd)

A distinct ED process may occur when both the controlling and dependent characters are of the BD type. Let us consider the evolution of tail armor in species with no tails, tails without armor, and tails with armor. Suppose the tail character evolves as before with multiple gains and losses. Obviously, gains of the tail armor can only occur if the tail is present. However, the evolution of the armor may differ from that of the tail color in the previous example because the birth of the tail may not necessarily imply the immediate emergence of the armor. It is biologically reasonable to assume a time lag between the births of these anatomical structures. Macroevolutionary lags are a common phenomenon (Erwin, 2021), for example, tetrapod limbs are anatomically dependent on the presence of bone tissue whose origin significantly predates that of the limbs (Niedźwiedzki et al., 2010; Keating et al., 2018). The lags between the births of characters specify the gradual emergence of anatomical hierarchies; not accounting for them makes a strong assumption that complex hierarchies may emerge over a short evolutionary time. For example, consider a multi-level hierarchy where an anatomical structure *S*_1_ controls another structure *S*_2_ that controls *S*_3_, and so forth til *S_n_*. If the lag condition is ignored, this hierarchy can instantaneously evolve into *S_n_* from *S*_1_. In other words, not correcting for the lags implies that a unicellular organism could evolve into an elephant over one evolutionary step, which is unrealistic. Thus, I find it essential to consider the ED process with lags separately and refer to it by the type of dependent character as the “birth-death ED process” (ED-bd). One possible realization for this process is given in Fig. 1B; it differs from the ED-ql realization by the lag following the second birth (*b*_2_ in Fig. 1B) of the tail. However, if one believes that a complex hierarchy can evolve instantaneously, the lag assumption should be removed, which would collapse the ED-bd to ED-ql.

In the probabilistic language, both the tail and armor characters can be viewed as two MCs that evolve similar to those from the tail color scenario, but the tail armor MC is permanently restricted to begin its evolution from the state “absent” to account for the lag.

### Modeling ED processes

As demonstrated above, it is natural to represent the simple ED processes by a duet of interacting Markov chains in which the evolution of one chain is embedded with a particular state of another. This modeling design is implemented in MSSD approach of Alekseyenko et al. (2008). In contrast to the described ED processes, MSSD model allows only one birth of the controlling character (e.g., tail), with the possibility of its subsequent losses in multiple lineages but without any secondary origins of this character. The stochastic realization for the MSSD process can be visualized by taking that for the ED-ql process (Fig. 1A) and prohibiting the second birth of the tail (*b*_2_ in Fig. 1A), including all the following transitions in the tail and color characters associated with that birth.

Another approach, SMMs with hidden states, models the ED process as the joint evolution of interacting Markov chains but does not explicitly account for the chain embedding; SMMs simulate the embedding using hidden states. Thus, both available models, MSSD and SMM, are different from the described ED processes. The following section describes the probabilistic models underlying the ED-ql and ED-bd processes for phylogenetic inference.

## Deriving Embedded Dependency Models

Below, I begin with deriving rate matrices for general ED models, then demonstrate how to model complex hierarchies and consider possible parametrizations of the new models for phylogenetic inference with morphological data.

As shown above, a simple ED process can be represented by a duet of MCs evolving in correlation. Thus, such a dependency should be appropriately modeled. Most Markov models in phylogenetics treat characters as independent entities and, thereby, cannot be applied to ED. The study of Goloboff et al. (2021) has recognized this fact that possibly led the authors to conclude that probabilistic treatment of ED requires new algorithms for likelihood calculation. They suggested that ED cannot be described with infinitesimal rate matrices (*Q*) alone, used for specifying MC evolution. Below, I demonstrate that this description is possible, using the SMM techniques for character amalgamation and aggregation (Tarasov, 2019, 2020). The representation of an ED model using a single rate matrix allows employing existing software and methods, such as matrix exponentiation and Felsenstein’s pruning algorithm (Felsenstein, 1981), for likelihood computation.

The general idea behind the derivation of ED models resides in combining two or more individual MCs into one by amalgamation. The amalgamated MC describes the joint evolution of the individual chains. Thus, one can use a single amalgamated MC instead of modeling the set of dependent chains to describe the same process. There are several types of amalgamations. For example, MCs can be amalgamated as independently evolving [the equation (2) in Tarasov (2019)] that is equivalent to the independent evolution of individual MCs. Hereafter, I refer to this amalgamation as “SMM amalgamation”. Its state space represents all possible state combinations from individual MCs. For example, if there are two binary MCs with the states {0,1} then their amalgamated chain includes four states {00,01,10,11}. In this paper, I introduce a new type of amalgamation, called the “ED amalgamation”, to construct rate matrices for ED processes. It is similar but different from the SMM amalgamation by the state space and rate matrix structure. The ED states are also combinations of the original states, but some redundant states are removed from *Q*. At the end of this section, I demonstrate how to use the combination of ED and SMM amalgamations to model custom character hierarchy.

### General ED models

In deriving the general ED models, I assume that all components of MCs (rates and initial vectors) might be different; in other words, these general models have the maximum number of possible parameters. The ED models with fewer parameters can be easily constructed from the general ones by linking the parameters. The mathematical notations below use bold font for vectors and capitals for matrices.

#### Qualitative ED model (ED-ql)

First, let us represent the tail and color characters from the described ED-ql process (Fig. 1A) as individual MCs and then derive their amalgamation. Naturally, each such an MC has two states that for brevity I denote as *T* – tail presence {*absent(a),present(p)*}, and *C* – tail color {*red(r),blue(b)*}. We assume that *C* and *T* evolve under the ED-ql process according to the initial probability vectors ***π_T_*** = (*π*_*T*_1__, *π*_*T*_2__) and *π*_*C*_ = (*π*_*C*_1__, *π*_*C*_2__), and these rate matrices:

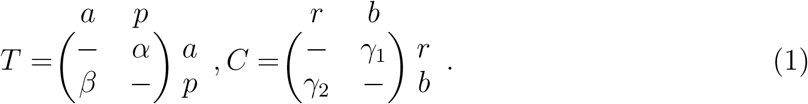

The MC for *T* characterizes the tail birth-death, while the embedded one for *C* specifies color state transitions. The amalgamated rate matrix for *T* and *C* that describes the ED-ql process is:

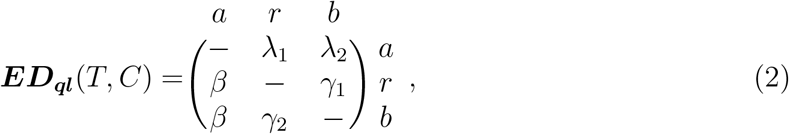

this matrix has three states {*a,r,b*} and the initial vector = (1 – *π*_*T*_2__, *π*_*T*_2__ *π*_*C*_1__, *π*_*T*_2__*π*_*C*_2__). Its rates λ_1_ and λ_2_ consist of two components: the transition rate *α* from *T*, and the embedded vector *ϕ_C_* = (*ϕ*_*C*1_, *ϕ*_*C*2_), so that λ_1_ = *ϕ*_*C*1_*α* and λ_2_ = *ϕC*_2_*α*. The embedded vector *ϕC* specifies the initial probabilities for the states in C when this character is allowed to evolve (i.e., when transiting to “tail” from “no tail”). I refer to this operation of combining T and C as the “ED-ql amalgamation”.

This amalgamation describes precisely the same ED-ql process shown in Fig. 1A; the proof of this is given in Appendix 1. Instead of the two individual MCs (T and C) with a total of four states, it uses one amalgamated MC with three states (Fig. 1C-E); this number is obtained by removing the redundant states. The amalgamated MC has a specific rate matrix; the maximum ED-ql model can have as many as five parameters where the same parameter denotes the transitions *r* → *a* and *b* → *a*. These constraints are important; relaxing them (e.g., making all rates different) would break the basic assumptions for the ED-ql process and indicate a complex state-dependent evolution between the controlling and dependent characters.

#### Birth-death ED model (ED-bd)

Now, let us consider the ED-bd process for the tail armor case (Fig. 1B). It can be characterized by two rate matrices, where the matrix for the tail is the same as *T* in the equation (1), and the matrix *A* – armor presence {*absent*(*a*_1_), *present*(*p*_1_)} denotes the birth-death of the armor:

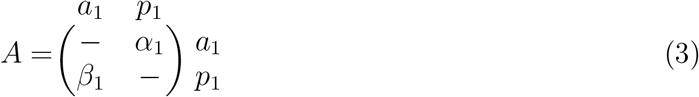

The amalgamation for the two BD characters *T* and *A* (hereafter, the “ED-bd amalgamation”) is different from the ED-ql amalgamation because the embedded MC for *A* should obey the gradual development of anatomical hierarchy. This yields the following amalgamated matrix:

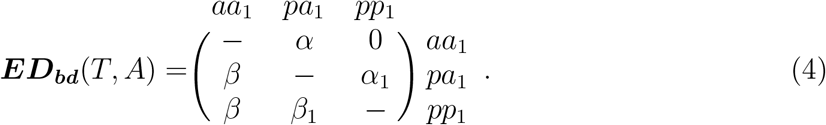

This matrix is similar to the ED-ql in the equation (2), but the embedded vector of the initial probabilities *ϕ_A_* for *A* is *ϕ_A_* = (1, 0); it penalizes the direct change from “no tail” (*aa*_1_) to “armor and tail present” (*pp*_1_) and constraints A to start its evolution from the state “tail present, no armor” (*pa*_1_). There are the following relationships between this matrix and the λ’s in the equation (2): λ_1_ = α_1_, λ_2_ = 0.

This amalgamation describes the same ED-bd process shown in Fig. 2A; the proof of this is given in Appendix 1. Like, the ED-ql, its state space is reduced to three states (Fig. 1D-F). The maximum ED-bd model can have as many as four parameters, with the same parameter denoting the transitions *pa*_1_ → *aa*_1_ and *pp*_1_ → *aa*_1_; the transition *aa*_1_ → *pp*_1_ is prohibited. Relaxing those parameters would break the basic assumptions for the ED-bd process.

**Fig. 2.**
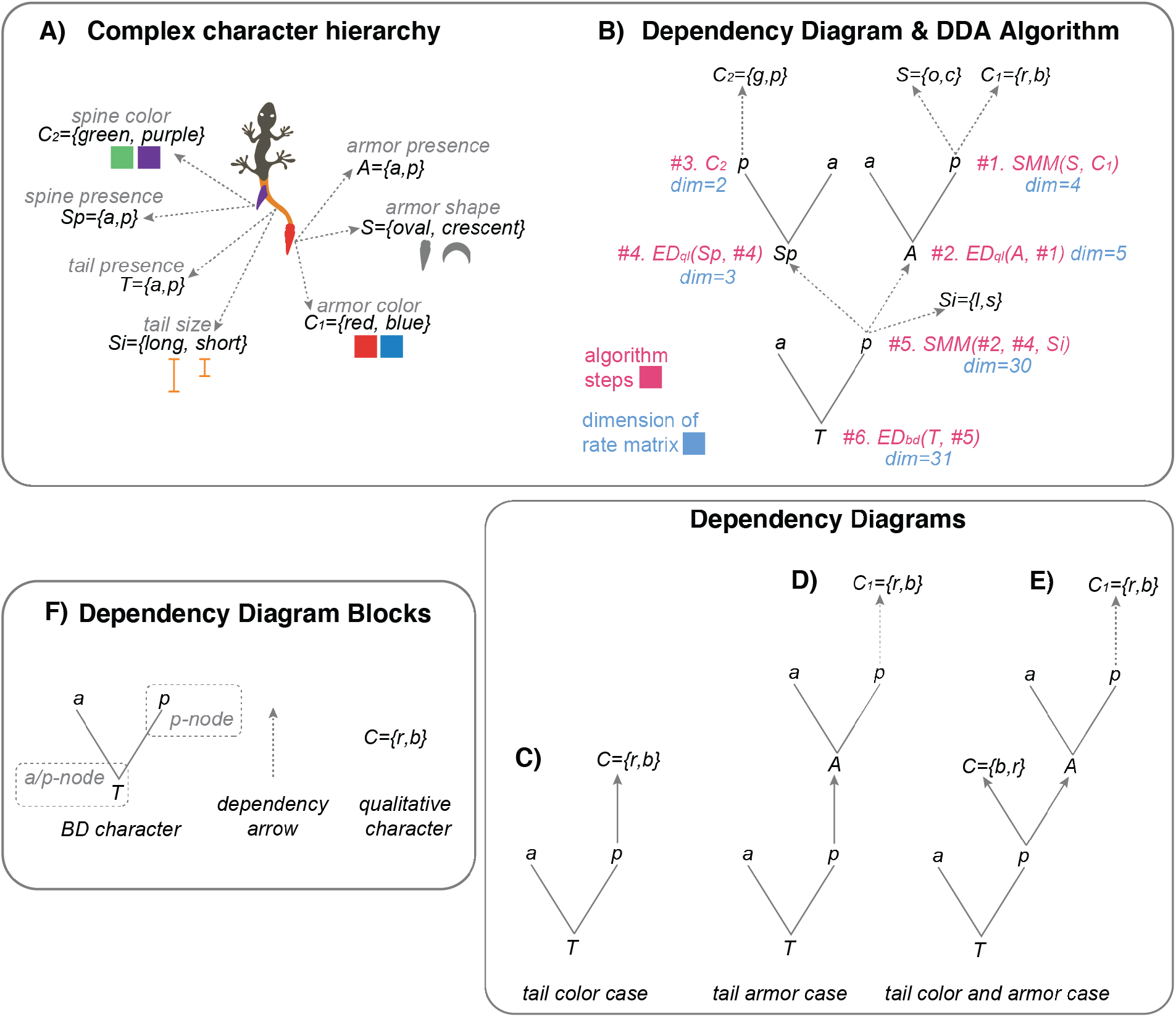
The dependency diagram amalgamation (DDA) algorithm. A) Complex character hierarchy for the hypothetical tail traits; it consists of three BD and four qualitative characters. B) The dependency diagram for the characters in (A); red color indicates the steps in the DDA algorithm, while blue color refers to the dimension of the rate matrices; *ED_ql_*, *ED_ap_, SMM* refer to the respective amalgamations. C-E) The dependency diagrams for the discussed character hierarchies. F) The building blocks of a dependency diagram.

#### Mixture of ED-ql and ED-bd

The cases when ED-bd and ED-ql processes co-occur in organisms are also common. The toy example of this is “the tail color and armor” (TCA) case. In which species have a two-level hierarchy: a tail with blue or red armor, a tail without armor, and no tail. To construct the rate matrix for TCA, one should successively repeat the ED-ql and ED-bd amalgamations:

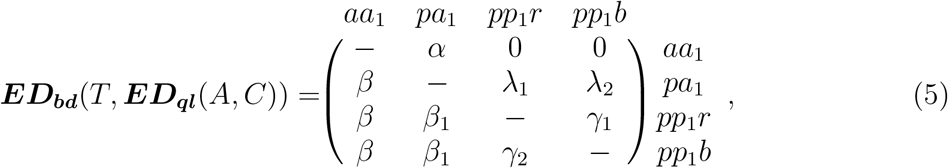

this matrix has the initial vector 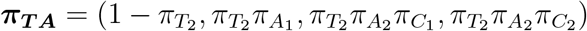; its λ_1_ + λ_2_ equals α_1_ from the equation (4). The maximum model has as many as seven parameters corresponding to interplay between the ED-ql and ED-bd processes.

#### General formula for ED

Suppose the controlling character (*Q_c_*) is the same as *T* in the equation (1), while the dependent character (*Q_d_*) may have any number of states and custom parametrization of the rate matrix. In that case, the general equation for the ED amalgamation is:

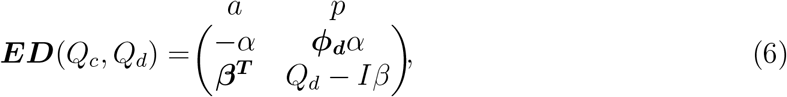

where *I* is the identity matrix, ***β*** is the vector of *β* rates, superscript ***T*** is an array transpose, and *ϕ_d_* is the appropriate type of the embedded vector for the initial probabilities of *Q_d_*. The amalgamated matrix’s dimensionality equals the number of states in *Q_d_* plus one. Note, this equation indicates that the type of ED solely depends on the embedded initial vector *ϕ_d_*.

#### Custom Character Hierarchy

Anatomical dependencies between body parts can generate character hierarchies of varying complexity. If a hierarchy includes several characters, it might be challenging to keep the right track on the sequence of character amalgamations. To facilitate the construction of amalgamated rate matrices, I propose Dependency Diagram Amalgamation (DDA) algorithm.

The DDA algorithm uses the dependency diagram (Fig. 2B-E) that schematically depicts relationships in character hierarchies and consists of three building blocks (Fig. 2F): (i) BD character; (ii) qualitative character; and (iii) dependency arrow. The BD character includes two subblocks: (i) *absence/presence (a/p) -node,* and (ii) *p – node,* which are the character itself and its state “present” (*p*), respectively. In the diagram, the BD and qualitative characters are interlinked through a dependency arrow to reflect hierarchical relationships. Specifically, the dependency arrow links the *p* – *node* of a controlling character with a dependent character (that can be either of the BD or qualitative type). The DDA algorithm traverses along the dependency diagram by amalgamating characters at its nodes producing the required rate matrix at the end. The ED characters are combined via ED-bd or ED-ql amalgamation, and independent characters are amalgamated via SMM.

For example, consider a complex hierarchy of three BD and four qualitative tail characters shown in Fig. 2A, for which we wish to construct a rate matrix. The DDA algorithm works as follows: it takes the required dependency diagram (Fig. 2B) and traverses it from the top to the root in a topologically ordered manner. At each *a/p*- or *p* – *node*, it combines all child characters into one matrix via SMM, ED-ql, or ED-bd amalgamations; the type of amalgamation depends on the node and its children (see Appendix 2). The amalgamation at the root of the diagram yields the final rate matrix. For our complex hierarchy, the diagram traversal requires six steps and returns the final matrix of dimension=31 (Fig. 2B).

The dependency diagram for the toy examples of tail color, tail armor, and TCA cases are shown in Fig. 2C-E. The vignette with the DDA algorithm for all considered examples is given in the package *rphenoscate*.

### Phylogenetic inference with ED models

The general ED models from the previous section can infer ancestral character states on a known topology. However, if the topology is the focus of analysis, they should be appropriately parameterized to avoid overfitting. For example, the ED-ql for the tail color case [the equation (2)] can have as many as five parameters that might be overwhelming given the limited number of data points in morphological datasets. I propose several parametrizations for ED models to balance model complexity and biological interpretation. These parametrizations use the toy examples of the tail color and TCA cases and can be further extrapolated to other EDs.

#### ED-ql model

The tail color ED-ql from the equation (2) converts into the Mk-like model (Lewis, 2001) by constraining it to have one free parameter (that is usually interpreted as a branch length), which gives the following rate matrix:

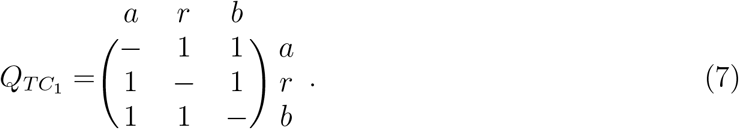

It is reasonable to assume that the transitions for tail gain, loss, and changes between the colors may occur at different rates. Thus, unlinking these parameters results in another model:

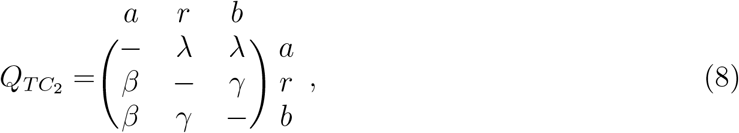

Constraining the sum of the rates to one as *β* + λ + *γ* =1 gives *γ* =1 – *β* – λ and reduces the total number of the parameters to two in this model.

#### Mixture of ED-ql and ED-bd

Unlike above, the TCA case from the equation (5), might be converted into the Mk-like model in two different ways because its rate matrix contains zero elements. The first type of the Mk parametrization makes all non-zero rates to be equal:

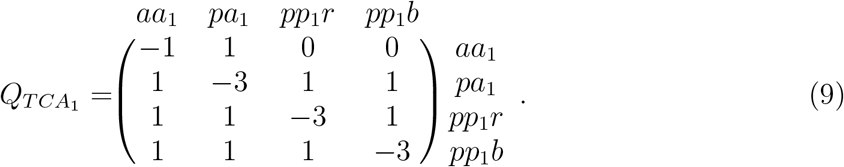

The second type of the Mk parametrization makes the main diagonal elements to be equal, while the off-diagonal rates become uniformly rescaled within each row:

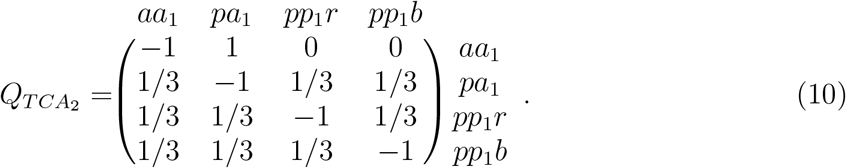

Although the models in the equations (9) and (10) have the same number of parameters (one), they may have different interpretations. The model *Q*_*TCA*_2__ assumes an equal expected waiting time to be spent in each state before leaving it, while *Q*_*TCA*_1__ assumes a longer time to be spent in the state *aa***i** than in the three other states. Thus, these models imply different evolutionary dynamics for the “no tail, no armor” state (*aa*_i_). The model *Q*_*TCA*_2__ might be be biologically more reasonable than Q_TCA1_ if there is no prior evidence to believe why the absence of the tail should evolve slower than its presence.

The third way to parametrize the TCA model is to assign its rates into different categories, which are of three main types – the rates responsible for the growth and loss of the hierarchy and changes between the qualitative states:

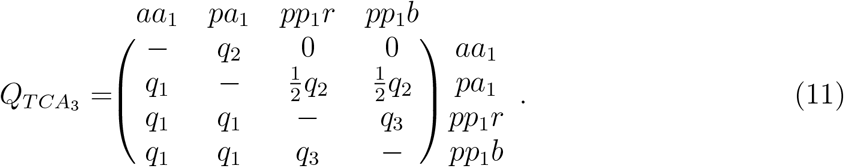

Constraining the sum of the rates to one as *q*_3_ = 1 – *q*_1_ – *q*_2_ reduces the total number of the parameters to two in this model.

Three types of initial vectors apply, in practice, to the proposed models: equal, equilibrium, and inferred state frequencies at the root of a tree. Note that all proposed models are generally time-irreversible due to their asymmetries in rates or the presence of zero rates. The time-irreversibility means that their likelihood should be calculated on a rooted tree (Felsenstein, 1981; Yang et al., 2006; Boussau and Gouy, 2006). In this regard, Bayesian dating analysis represents a convenient framework for their implementation. The only model in this set that is trivially time-reversible is *Q*_*TC*_1__ when its initial vector has equal or equilibrium state frequencies (both are the same). Thus, the phylogenetic inference with this model does not require a rooted tree – the setting that is standard to many maximum-likelihood programs.

## Materials and Methods

### Availability

The functions and vignette for the DDA algorithm are provided in the R package *rphenoscate* https://github.com/uyedaj/rphenoscate [To be submitted to CRAN soon]. The scripts to reproduce the simulations below are available at GitHub repository https://github.com/sergeitarasov/MorphoModels.

### Simulations

To assess the ED models, I performed three series of simulations in the Bayesian framework using RevBayes: (1) the first series tests the topological and model performance between SMMs and various parametrizations proposed for the ED models using fixed data; (2) the second series demonstrates that the ED models and SMMs may yield different topologies using simulated data; and (3) the third series tests the behavior of the ED models in the original formulation of the tail color problem that is commonly used for assessing inapplicable characters (Maddison, 1993; Hawkins et al., 1997). Further details on these three series of simulations are given below.

Since, generally, the ED models are time irreversible and require a rooted tree, I performed all analyses in the dating framework using the global molecular clock model (Zuckerkandl and Pauling, 1962). I used the branch rate prior *ψ* ~ *Exponential*(10), Yule prior distribution for the tree topologies, and the root age set to 1.0. The likelihoods were conditioned on observing only variable characters in the data (Lewis, 2001). Given the simplicity of my toy datasets, I use 200-500K generations in the Markov Chain Monte Carlo algorithm and two independent runs for each simulation.

The relative model performance was evaluated using Bayes factors (BF) which compares the ratio of marginal likelihoods (*Mln*) of two candidate models (estimated as *Mln* of the best model minus *Mln* of the focal model). The marginal likelihood was calculated using the stepping-stone algorithm (Xie et al., 2011) implemented in RevBayes by averaging over the two runs.

#### 1. ED models and SMMs: fixed data

This series compares eight different ED models (ED1-ED8) against themselves and two SMMs (Mk-SMM-ind and Mk-SMM-sw). I use the dataset of four species with fifty identical four-state characters referring to the TCA case (Fig. 2E) to perform tree inference and calculate the marginal likelihood. This dataset was used by me earlier to assess SMMs and was shown to contain sufficient information content for model assessment (Tarasov, 2019).

The models ED1-ED8 (Table 1) were constructed as combinations of the three rate matrices *Q*_*TCA*_1__, *Q*_*TCA*2_ and *Q*_*TCA*_3__ from the equations (9–11) and the three types of the initial vectors at tree root with equal, equilibrium and inferred probabilities. The tested models range in the number of parameters from one to four. This number refers exclusively to the components of the Markov model and calculates as the sum of free parameters in *Q* + *π* + 1 (one stands for the branch rate *ψ*). The combinations of *Q*_*TCA*_3__ and the inferred initial vector were not included in the simulations due to the excessive number of parameters in them.

**Table 1.** The Performance of the ED models under fixed data. The “Topology” column indicates the tree A or B from Fig. 3.

| Model | Parameters | Marginal Ln | BF | Topology |
| --- | --- | --- | --- | --- |
| The ED models: |  |  |  |  |
| 1 ED5 ($Q_{TCA_1}, \pi = \text{inferred}, \psi$) | 4 | -292.29 | 0.00 | A |
| 2 ED3 ($Q_{TCA_1}, \pi = \text{equal}, \psi$) | 1 | -296.13 | 3.85 | A |
| 3 ED8 ($Q_{TCA_2}, \pi = \text{inferred}, \psi$) | 4 | -299.10 | 6.82 | B |
| 4 ED4 ($Q_{TCA_1}, \pi = \text{equilibrium}, \psi$) | 1 | -300.50 | 8.22 | A |
| 5 ED6 ($Q_{TCA_2}, \pi = \text{equal}, \psi$) | 1 | -303.65 | 11.36 | B |
| 6 ED7 ($Q_{TCA_2}, \pi = \text{equilibrium}, \psi$) | 1 | -304.98 | 12.69 | B |
| 7 ED1 ($Q_{TCA_3}, \pi = \text{equal}, \psi$) | 3 | -308.24 | 15.95 | A |
| 8 ED2 ($Q_{TCA_3}, \pi = \text{equilibrium}, \psi$) | 3 | -313.42 | 21.13 | B |
| The expanded ED models and SMMs: |  |  |  |  |
| 1 ED3 ($Q_{TCA_1}, \pi = \text{equal}, \psi$) | 1 | -298.55 | 0.00 | A |
| 2 ED6 ($Q_{TCA_2}, \pi = \text{equal}, \psi$) | 1 | -305.14 | 6.60 | B |
| 3 Mk-SMM-sw ($\psi$) | 1 | -318.55 | 20.00 | A |
| 4 Mk-SMM-ind ($\psi$) | 1 | -323.38 | 24.84 | A |

The models Mk-SMM-ind and Mk-SMM-sw for TCA have one parameter, eight hidden, and four observable states [page 3 in Tarasov (2019)]. In applying these models, I use a uniform vector of initial probabilities for the observable states. The conventional representation of ED models for TCA has four observable and no hidden states. The state space of ED models can be expanded with hidden states to match the SMMs. This expansion might help assess model behavior (Tarasov, 2019); it creates an eight-state rate matrix that, if lumped, collapses back to the original four-state one. The expanded and original models are congruent, meaning they return the same likelihood values; however, their posterior probabilities may differ due to the parametrization in priors. Generally, the congruency indicates that the original four-state ED and eight-state Mk-SMM models can be compared using marginal likelihood regardless of differences in their state spaces. However, I use both the expanded and original ED models for demonstrative purposes in this study.

#### 2. ED models vs. SMMs: simulated data

The series of simulations aims at showing that the ED models may behave differently from SMMs by recovering distinct topologies. I simulated 1000 trees in *phytools* (Revell, 2012) under the Yule process with a birth rate 0.1. Then, I simulated ten variable characters, corresponding to the TCA case, under the ED6 model (*Q*_*TCA*_2__) for each tree. Next, I estimated trees under four models – the original one, ED3, Mk-SMM-ind and Mk-SMM-sw – for each simulated dataset. The Robinson-Foulds distance [RF, Robinson and Foulds (1981)] between the true tree and the obtained majority rule (50%) consensus (MJ) tree was calculated for assessing the topological accuracy (Wright and Hillis, 2014).

To compare the proportions of correctly recovered trees between the original and other models (i.e., RF=0), I used the proportion differential (pd) and bootstrapped p-values. The proportion differential calculates as *pd* = *Prop*(*ED*_6_) – *Prop*(*M_i_*), where *Prop* stands for the proportion of the correctly recovered tress, and *M_i_* is one of the models. The p-value for *pd* was estimated by bootstrapping with 5000 replications. This p-value shows if the proportion of correctly recovered trees is statistically significant between the models.

#### 3. ED and Tail Color Problem

In this series of simulations, I test if the ED models can appropriately model the tail color problem (TCP). I follow the same simulation setup as before (Tarasov, 2019). The TCP considers a tree of 14 species where the tailed species (with red and blue tails) are nested within the left and right clades of the tailless species (Maddison, 1993). The tree (Fig. 3C) is assumed to be fully resolved except for the relationships in the left-tailed clade (LTC); all resolved clades are supported by at least one binary character to avoid zero-length branches. The tail color is encoded as the three-state character (no tail, tail red, tail blue) and replicated fifty times to provide sufficient information for tree inference. The goal of these simulations is to assess possible resolutions of the LTC. For this purpose, I use one-parameter (*Q*_*TC*_1__, the equation 7) and three-parameter (*Q*_*TC*_2__, the equation 8) with equal state frequencies at the tree root. For each of these models, I performed two simulations with a varying number of synapomorphies supporting the LTC (one or eleven characters) to assess the effect of branch length on the topology.

**Fig. 3.**
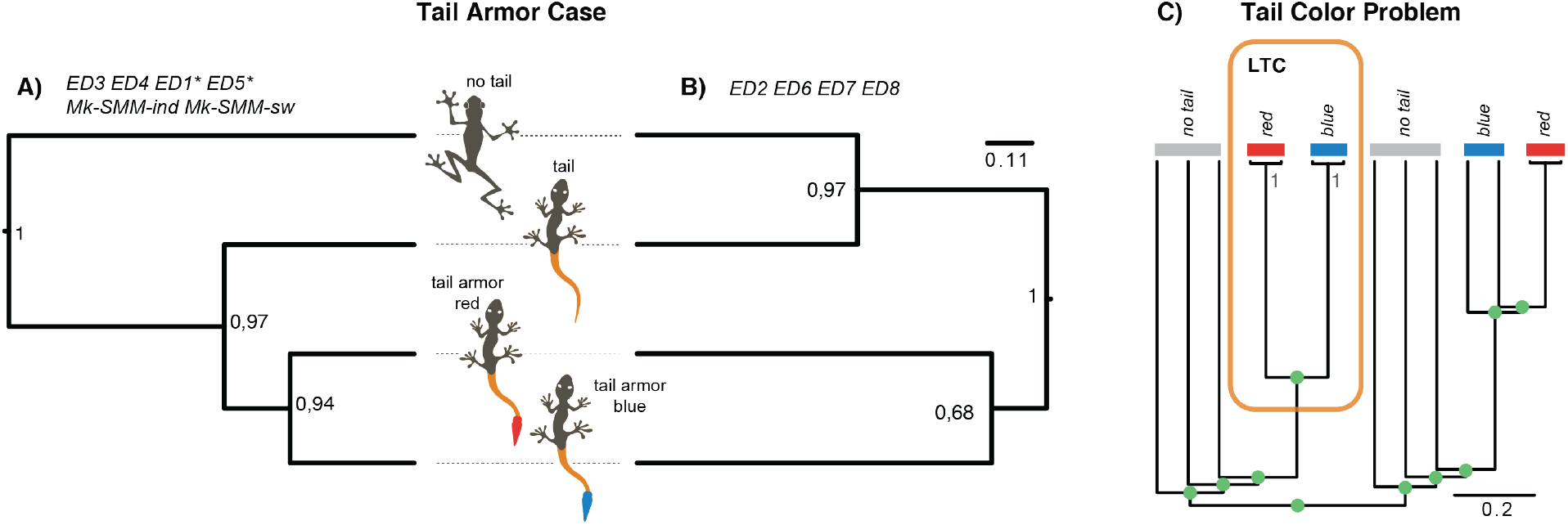
The assessment of the ED models. A-B) The two majority rule (50%) consensus trees, which were recovered by eight ED models and two SMMs for the TCA case; the values at tree nodes indicate posterior probabilities from the analyses with ED4 (A) and ED8 (B) models (they are selected for the demonstrative purpose only); “*” indicates that “tail” and “no tail” taxa collapse into a polytomy on the majority rule (50%) consensus trees but are resolved in MCC tree that is the same as the tree in (A). C) The tree that exemplifies the tail color problem where the tailed species are nested within the two major clades of the tailless species; all relationships are assumed to be resolved (green balls) except those in the left tailed clade (LTC) that includes two species with a red tail and two species with blue tail respectively; the values in the LCT indicate posterior probabilities.

## Results

### 1. ED models and SMMs: fixed data

The ED models (Table 1) produce two distinct tree topologies for the TCA case when the data are fixed (the fifty identical characters). One topology suggests a gradual appearance of the tail and tail armor from no tail state (Fig. 3A); it is recovered by the models ED1, ED3-5 on the maximum clade credibility (MCC) tree. The MJ tree for these models is the same as the MCC one, except for ED1,5 where “tail” and “no tail” taxa collapse into a polytomy. Another topology groups together species with and without the armor (Fig. 3B); it is recovered by the models ED2, ED6-8 on both the MCC and MJ trees. Both SMMs (Mk-SMM-sw and Mk-SMM-ind) yield the same tree that reflects the gradual evolution of the tail and armor (Fig. 3A).

The marginal likelihood values significantly vary across the ED models (Table 1). The best fit models are ED5 and ED3, which reflect the gradual appearance of the tail from no tail (Fig. 3A). These two models have the Mk-like rate matrix with equal transition rates (*Q*_*TCA*_1__, the equation 9). The worst performing models are ED1 and ED2 with the two-parameter rate matrix (*Q*_*TCA*_3__, the equation 11). The *BF* value between the best fit model (ED5) and the worst fit model (ED2) is 21.13, which points to a substantial overperformance of ED5 over ED2 in these toy simulations.

I selected two simple one-parameter models, ED3 and ED6, to assess their fit compared to one-parameter SMMs for inapplicable characters (Mk-SMM-ind and Mk-SMM-sw); the state space of ED3 and ED6 was appropriately expanded with hidden states. The selected ED models performed decisively better than the respective SMMs in these toy simulations (*BF* >> 10, Table 1).

### 2. ED models vs. SMMs: simulated data

The ED models and SMMs behave distinctly in recovering the correct tree using simulated data. The true tree supported by ten characters, simulated with the ED6 model, was correctly recovered in 54%, 53%, 51%, and 51% simulations under ED6, ED3, Mk-SMM-ind and Mk-SMM-sw models, respectively. Although these proportions are similar across the tested models, the estimated p-values support the statistically higher rate of the true tree recovery for the ED6 compared to Mk-SMM-ind and Mk-SMM-sw models (p-values: 0.0042 and 0.0036, respectively). There was no statistically significant difference between the performance of ED6 and ED3 models (p-value 0.19).

### 3. ED and Tail Color Problem

The ED models show the appropriate behavior for the TCP. Both *Q*_*TC*_1__ and *Q*_*TC*_2__ correctly resolve the LTC – they support the monophyly of the red- and blue-tailed species with the posterior probability 1.0 (Fig. 3C). The same topology is recovered regardless of the number of synapomorphies (one or eleven) used in the simulations.

## Discussion

### Assessing the proposed ED models using the simulations

Coding the tail color as inapplicables, in the context of the TCP, was shown to suffer from the undesirable behavior in parsimony analysis by incorrectly recovering the species relationships within the LTC (Maddison, 1993). Previously, I showed that, unlike parsimony, SMMs (e.g., Mk-SMM-ind and Mk-SMM-sw) provide the solution to modeling the inapplicables (Tarasov, 2019). In this study, I have demonstrated that the ED models, like SMMs, also solve this problem – they correctly resolve the LTC and accommodate dependencies. Thus, the three-state Mk model [the equation (7)] does provide appropriate treatment for the simple ED-ql process. This result contradicts my previous conclusion in Tarasov (2019) that now should be considered incorrect; the Mk model can, in fact, model anatomical dependencies. However, it is only applicable to a simple ED-ql process but not the ED-bd or complex hierarchies, which should be handled using the appropriate amalgamations and DDA algorithm.

One can construct several ED models for a simple dataset, as shown for the tail color and TCA cases, which may differ in rate matrices and initial vectors. The first series of simulations with fixed data demonstrates that the fit and topology of different models can vary significantly due to their parametrizations that imply different assumptions for character evolution. Specifically, the ED models recover two topologies – one assuming gradual evolution of the tail and tail armor, and the other grouping “tail”, and “no tail” species (Fig. 3A-B). This result tends to be associated with the parametrization of the rate matrix: the matrix *Q*_*TCA*_1__ with equal transition rates usually produces the tree in Fig. 3A, while the matrix *Q*_*TCA*_2__ with equal diagonal elements recovers the tree in Fig. 3B.

In the simulations with the fixed data, SMMs support only one tree, while the ED models yield two trees. In the simulations with simulated data under the ED6 model, the proportion of correctly recovered trees by SMM was statistically lower than by the ED models. Thus, these results suggest that the ED models and SMMs are similar but may behave differently and recover different trees depending on the dataset.

The parametrization of *Q* and the initial vector can affect model marginal likelihood. In present simulations, the ED models outperform SMMs; the best-fit ED model (ED5 with four parameters) comes to support the gradual evolution of the tail and armor. However, this result may differ depending on the dataset. In practice, one should always compare the ED models and SMMs using statistical methods for model selection (e.g., Akaike information criterion, BF), as shown in the first series of simulations.

The proposed ED models are also different from the stochastic Dollo model (MSSD) because of their different assumptions for the number of births in the controlling character. The available implementations of MSSD in BEAST (Drummond et al., 2012) and RevBayes (Tarver et al., 2018) can handle only a one-level hierarchy in which the controlling character embeds the evolution of one or more dependent ones. Because of this limitation, the MSSD models were not tested in the present simulations since they do not work with the repetitive births in the controlling character (TCP) and a two-level anatomical hierarchy (TCA case). However, one should expect that the performance of the ED and MSSD should be markedly different due to their assumptions.

### Modeling Character Hierarchies

The ED exhibits a common phenomenon in morphology since organismal body parts often depend on each other. It occurs when one character is allowed to evolve only within a particular state of another character. The ED has often been coded using inapplicable characters, which are then analyzed under parsimony or likelihood-based methods. In parsimony, the inapplicable coding was shown to be inappropriate (Maddison, 1993), which encouraged the development of new parsimony-based methods to handle the ED (Brazeau et al., 2019; Hopkins and St. John, 2021; Goloboff et al., 2021).

In this paper, I used theoretical derivations to show that specific Markov models can be straightforwardly adopted to model the ED explicitly. These models are defined using single rate matrices via the appropriate amalgamation of the involved characters [the equation (6)]. They do not require unique algorithms for likelihood computations and run in any software that allows custom specification of *Q*’s and inference on rooted topologies (since many models are time-irreversible). Specifically, I demonstrated how the Bayesian tree inference with the new models could be conducted in RevBayes. The ED models come in two flavors – ED-ql and ED-bd – depending on the type of character dependency. They differ from SMMs, which do not explicitly imply the ED but model it using hidden states. Note that the performance of the ED models and SMMs is data-dependent, and the best model for a specific character hierarchy should be selected using model selection criteria.

The probabilistic phylogenetics has only a limited set of models available for discrete traits (Lewis, 2001; Klopfstein et al., 2015; Pyron, 2017; Felsenstein, 2005; Wright et al., 2016). This limitation imposes certain constraints for tree inference when trying to account for a diverse array of processes that generate trait evolution. The new ED models expand the current arsenal of modeling approaches for morphology. The flexibility of ED models and SMM compared to parsimony lies in their natural ability to model the trait-generating process and, thereby, dependencies. The combination of ED models and SMMs can potentially account for any dependency that may exist between traits.

### Practical Recommendations

In practice, anatomical dependencies between characters may be complex. To use the ED models in phylogenetic inference, first, one must construct the amalgamated rate matrices using the DDA algorithm and, for instance, the package *rphenoscate*. Each separate hierarchy can have its own amalgamated rate matrix. Next, the characters in the data matrix should be appropriately recoded. The character hierarchies are usually coded using a set of characters. For example, the tail color case often uses two characters (one coded as inapplicable), while the TCA uses three characters (two coded as inapplicables). Note that all the ED models require representing these sets of characters as one single character whose states must match those in the constructed ED rate matrix. Thus, the characters defining a hierarchy should be recoded into one character before running an analysis. This procedure is partially automatized in the package *rphenoscate*.

The recoding results in a set of characters, in which each character represents a separate hierarchy. All of them can be placed in a separate partition subset to which a selected ED model is assigned. The ED models vary in the number of parameters (e.g., from one to two for the simple models). The models with the unlinked parameters [the equations (8) and (11)] can be applied to model rate heterogeneity between different types of transitions – gain and loss in the BD character(s) and transitions within the qualitative characters. Additionally, the rate heterogeneity across the characters, if there are several of them (each corresponding to a separate hierarchy), can be modeled by adopting the discrete-gamma model (Yang, 1994). The likelihoods of the ED models should be conditioned on observing variable characters only (Lewis, 2001) since constant characters are not sampled in morphological datasets.

The proposed approach for constructing ED models cannot be practically applied to molecular data, for example, for simultaneous modeling of the gene’s birth-death process and its nucleotide substitution. Although the required ED model can be constructed, its dimensionality would be computationally prohibitive. Deriving such a model requires SMM amalgamation for all individual sites and ED-bd amalgamation for the gene’s birth-death process. The number of states in such an amalgamated rate matrix is 4^*N*^ + 1, where *N* denotes the number of gene sites; this number is enormous even for a locus with hundred sites, which prevents computing the matrix exponential. Thus, another way of correcting Markov model embedding in likelihood calculation should be developed for molecular data to avoid using large rate matrices. The proposed models are suitable for only moderate-size hierarchies occurring in organismal morphologies.

## Acknowledgements

I am thankful to Diego Sasso Porto and Josef Uyeda for discussing anatomical dependencies and helping integrate the relevant functions into the *rphenoscate* package.

## Appendix

## Appendix 1: Proofs for ED models

Herein, I prove that the amalgamated ED matrices in the equations (2) and (4) correspond to the ED-ql and ED-bd processes, respectively. This proof uses the techniques of state aggregation and lumpability of MCs (Tarasov, 2019).

### Lumpability

In a nutshell, two or more states of an initial MC can be aggregated (lumped) into one state that produces the aggregated process with fewer states. If the aggregated process is still Markovian, the initial MC is called lumpable for the given state aggregation. If the process is not Markovian, the initial aggregation is called non-lumpable. The lumpability property guarantees that the transition rates of the lumped MC can be derived from the initial chain and thereby modeled using the lumped MC. There are several classes of lumpability, of which the “strong lumpability” is relevant to this paper due to its mathematical properties (Tarasov, 2019). Hereafter, I refer to the “strong lumpability” as simply “lumpability”. MC is lumpable under any value of the initial probability vector if certain rate symmetries in *Q* hold. These symmetries should obey the “row-wise sum rule” (RWR). Aggregation of states in a Markov chain can be seen as partitioning the transitions in the initial Q into blocks (subsets); these blocks are the transitions in the aggregated matrix. The RWR implies the following: the original MC is lumpable with respect to a given partitioning when the row-wise sum of rates within one partition block in the original *Q* is the same for all rows within the given partition block, and this property holds for all blocks in the matrix (Kemeny and Snell, 1960); the rates in the aggregated matrix represent simply row-wise sums of the original rates. For details, I suggest consulting my earlier work (Tarasov, 2019).

The lumpability property is general and applies to all MCs. Herein, this property is used to analyze dependencies between the controlling and dependent characters in an ED. If *X* and *Y* are MCs, and *X* is independent with respect to *Y*, then the amalgamated MC for *X* and *Y* should be lumpable with respect to *X*.

### Necessary and sufficient conditions

I propose that the ED implies the following necessary and sufficient conditions to hold: (1) the controlling character is independent of the dependent character; (2) the dependent character is dependent on the controlling one; (3) after a certain time, the probability of states in the dependent character depends only on the last realized state in the controlling character. The last condition indicates the embedded evolution of the dependent character when the controlling one is present. In the context of the tail color case it means that the probability of observing *C* (*r* or *b* state) at time *τ* (Fig. 1A) depends only on the last time interval *τ* – *t* of having the tail absent or present but not the previous tail occurrences, which is *Pr*(*C, τ*) = *Pr*(*C, τ* – *t*).

Hereafter, I call these conditions the “ED conditions”. They apply to all types of ED processes. Proofing that the ED conditions hold in an amalgamated model implies that this model describes the ED process.

### ED-ql model

The ED-ql matrix from the equation (2) fulfills all three ED conditions given above. Note, it is lumpable with respect to the aggregation of states {*a*, {*r, b*}} because the transitions *r* → *a* and *b* → *a* have the same parameter, which maintains the RWR. This lumping yields the matrix *T* and indicates the independence of *T* from *C* since no information in *C* is needed to characterize *T* (condition 1). In turn, *C* has no lumpable aggregations since all possibilities break the RWR, indicating its dependency on *T* (condition 2). The proof of condition three is given below.

### ED-bd model

The ED-bd matrix from the equation (4) fulfils all ED conditions. It is lumpable with respect to the aggregation of states {*aa*_1_, {*pa*_1_,*pp*_1_}} because *pa*_1_ → *aa*_1_ and *pp*_1_ → *aa*_1_ have the same parameter that maintains the RWR. This lumping yields the matrix *T* and indicates the independence of *T* from *A* since no information in *A* is needed to characterize *T* evolution (condition 1). In turn, *A* has no lumpable aggregations indicating its dependency on *T* (condition 2). The proof of condition three is given below. Note that the differences between the ED-bd and ED-ql models lie in the embedded dependency vector ***ϕ*** that penalizes the instantaneous growth of anatomical hierarchies.

An alternative proof can be constructed to show that ED matrices in the equations (2) and (4) describe the ED-ql and ED-bd processes. This proof can be developed by showing that the transitions implied by the ED-ql and ED-bd processes (Fig. 1A-B) are isomorphic to those in the equations (2) and (4). Obviously, it is true. There are six transitions for the ED-ql and four for the ED-bd process, which are the same as those in the equations (2) and (4), respectively.

### Proof of the Condition 3

This condition suggests the following equality to hold for the dependent character *Q_d_*, *Pr*(*Q_d_,τ*) = *Pr*(*Q_d_,τ* – *t*) meaning that at time *τ*, the evolution of *Q_d_* depends solely on the last time interval *τ* – *t* of having the controlling character *Q_c_* in a state absent or present but not the previous *Q_c_*’s states.

I will derive the joint distribution of the controlling *Q_c_* and dependent *Q_d_* characters using the two approaches: (i) the conditional approach 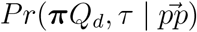 that directly implies the condition 3; and (ii) the direct calculation using, as an example, matrix exponential for the ED-ql model from the equation (2), i.e., 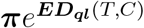 (the formula is not given here for brevity); the same logic applies to ED-bd model from the equation (4). The equality of the two approaches would suggest that condition 3 is true. For tractability, suppose that *Q_c_* is time-reversible:

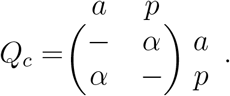

The probability of last *p* in time-reversible *Q_c_* is hence exponential

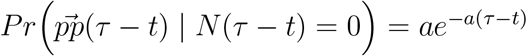. Herein, the notation 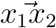 indicates a transition from the state *x*_1_ to the state *x*_2_; *N*(*t*) counts the number of transitions to different states. Two scenarios are possible:

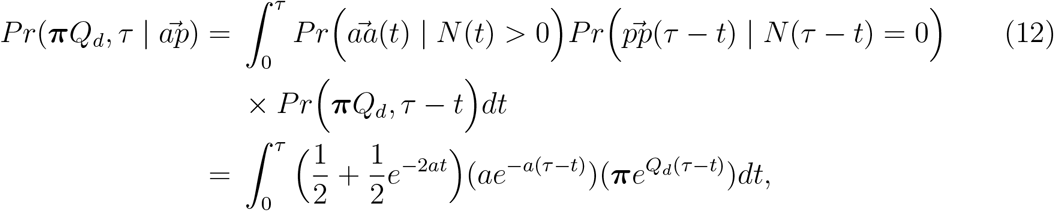

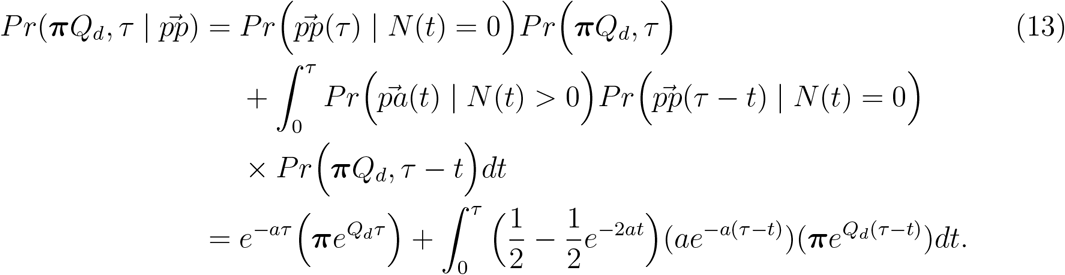

Now averaging over the initial vector:

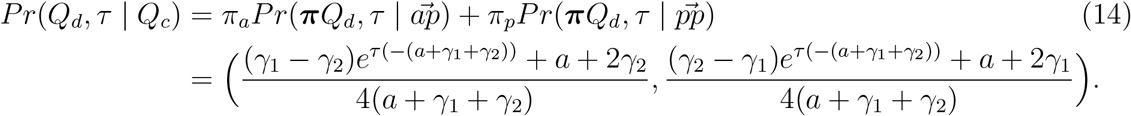

Note, *Pr*(*Q_d_,τ* | *Q_c_*) in the equation (14) is the same as 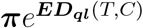 which proofs the aforementioned condition 3.

## Appendix 2: The Dependency Diagram Amalgamation (DDA) algorithm

As an input, the DDA algorithm requires a dependency diagram (e.e., Fig. 2D). The algorithm works by taking the diagram and traversing it from tips to root in a topologically ordered manner; the DDA amalgamates characters during the traversing. The amalgamations occur only at *a/p*– or *p* – *nodes* using the following rules:

1. At each *p* – *node* combine all child characters via SMM amalgamation [the equation (2) in Tarasov (2019)].
2. At each *a/p* – *node* conditionally combine all child characters via:

a. ED-ql amalgamation if the children are qualitative characters (e.g., as in the equation 2).
b. ED-bd amalgamation if the children are BD characters (e.g., as in the equation 4).
c. The mixture of ED-ql and ED-bd amalgamations if the children are BD and qualitative characters; this requires appropriate construction of *ϕ_d_* to match the dependencies between states (the equation 6).
3. The amalgamation at the root of the dependency diagram yields the desired final rate matrix.

